# High Recovery Stop-and-Go Extraction Tips Using Polytetrafluoroethylene Disks Embedded with Poly(styrene-divinylbenzene) Particles for Proteomics

**DOI:** 10.64898/2026.01.04.697531

**Authors:** Hiroto Kakiuchi, Eisuke Kanao, Kosuke Ogata, Yasushi Ishihama

**Affiliations:** Graduate School of Pharmaceutical Sciences, Kyoto University, Kyoto 606–8501, Japan; Laboratory of Proteomics for Drug Discovery, National Institute of Biomedical Innovation, Health and Nutrition, Ibaraki, Osaka 567–0085, Japan

**Author notes:** Correspondence to: Yasushi Ishihama, Graduate School of Pharmaceutical Sciences, Kyoto University, 46–29 Yoshidashimoadachi-cho, Sakyo-ku, Kyoto 606–8501, Japan.

**Keywords:** StageTip, desalting, proteomics, peptide purification, sample pretreatment

## Abstract

The stop-and-go extraction tip (StageTip) is widely used for peptide purification in bottom-up proteomics, yet the original 3M Empore^TM^ disk is no longer available, prompting the need to evaluate current alternatives. Two types of commercially available poly(styrene-divinylbenzene) (SDB)-containing polytetrafluoroethylene (PTFE) disks were used to fabricate StageTips. Desalting was performed on 500 ng and 20 ng of HeLa tryptic peptides, and the nanoLC/MS/MS results were compared. Both StageTips demonstrated equivalent performance for the 500 ng sample, but for the 20 ng sample, one StageTip identified 1.8 times more peptides, particularly longer and more hydrophobic peptides. Physical characterization and scanning electron microscope imaging of these disks revealed that the high-performing disk contains more PTFE fibers, partially covering the SDB particle surface. This likely prevents hydrophobic peptides from being trapped in small mesopores, improving recovery from low-input samples. These findings demonstrate that disk material properties critically influence performance in trace proteomics.

## 1. INTRODUCTION

Since its introduction in 2003, the stop-and-go extraction tip (StageTip) has become a widely used sample-preparation tool in LC/MS/MS-based bottom-up proteomics^1,2)^. A StageTip consists of chromatographic particles densely and thinly packed inside a pipette tip, enabling rapid loading, washing, and elution by centrifugation or manual force. Each step can be completed within seconds. In contrast, commercially available tip-based solid phase extraction (SPE) devices such as ZipTip, or self-packed loose-bed tips prepared from bulk particle suspensions^3,4)^ require sufficient equilibration time to allow mass transfer of peptides to the solid phase. As with conventional SPE, these formats generally do not achieve high recovery when operated under the fast conditions typical of StageTips^5,6)^. To address this limitation, alternative workflows have been proposed, such as adding TiO_2_ particles directly to peptide solutions, equilibrating by stirring for approximately 30 minutes, and subsequently loading the mixture onto a frit-containing pipette tip for phosphopeptide enrichment^7)^.

The original StageTip utilized 3M Empore^TM^ SDB-XC disks, in which 10-µm porous SDB particles (80 Å pore size) were densely embedded within a 0.5-mm thick PTFE sheet, forming 0.5 µm through-pore structure^8)^. These disks enabled quantitative recovery of 5 fmol to 75 pmol of BSA digest even at flow rates of 5 µL/s through a 0.4-mm diameter disk^1)^. However, production of this material was discontinued several years ago. In this study, we evaluated two currently available porous SDB-embedded PTFE disks from different manufacturers using tryptic digests of HeLa cell lysates. Given the substantial improvements in MS sensitivity over the past two decades and the increasing demand for low-input proteomics, we compared peptide recovery at two sample amounts—500 ng and 20 ng—to assess their suitability for modern micro- and nanoscale workflows.

## 2. EXPERIMENTAL

### 2.1. Materials

UltraPure Tris was purchased from Thermo Fisher Scientific (Waltham, MA, USA). Sequencing-grade modified trypsin was purchased from Promega (Madison, WI, USA). Water was purified by an ELGA PURELAB Quest 2 system (High Wycombe, UK). CDS Empore™ SDB-XC (CDS-SDB) disks were purchased from GL Sciences (Tokyo, Japan). Affinisep BioSPE^TM^ SDB (AFS-SDB) disks were purchased from Practical (Chiba, Japan). Protease inhibitors were purchased from Sigma-Aldrich (St. Louis, MO, USA). Blunt-end syringe needles were purchased from Hamilton (Reno, NV, USA) and 200 μL pipet tips were purchased from Gilson (Middleton, WI, USA) for the preparation of StageTips. All other chemicals and reagents were purchased from Fujifilm Wako (Osaka, Japan) unless otherwise specified.

### 2.2. Scanning electron microscope analysis

Surface morphology of the SDB disks was examined using a scanning electron microscope (SEM) JCM-7000 (JEOL, Tokyo, Japan) operated in high-vacuum mode. Disk samples were cut into approximately 5 mm pieces and mounted on aluminum pin stubs using carbon conductive tape. Surface observation was performed without additional coating or modification. Imaging was conducted with the BED-S detector at an accelerating voltage of 15.0 kV, working distance (WD) 11.7 mm, and High-PC beam current mode. The microscope’s control software was used to generate and export digital images.

### 2.3. Peptide sample preparation

HeLa S3 cells obtained from the JCRB Cell Bank (Osaka, Japan) were cultured to 80% confluency in Dulbecco’s modified Eagle’s medium containing 10% fetal bovine serum in 10 cm diameter dishes. Cells were washed twice with ice-cold PBS, collected using a cell scraper, and pelleted by centrifugation. Protein extraction and trypsin digestion was done by the phase-transfer surfactant (PTS) protocol as previously described^9)^. Briefly, pellets were lysed in 12 mM sodium deoxycholate / 12 mM sodium lauroyl sarcosinate / 100 mM Tris-HCl (pH 9.0) with protease inhibitors, heated at 95 °C for 5 min, sonicated 20 min, quantified by BCA assay, reduced with 10 mM DTT (30 min), alkylated with 50 mM IAA (30 min, dark), diluted 5× with 50 mM ammonium bicarbonate, and digested with Lys-C (3 hr, room temperature) followed by trypsin (overnight, 37 °C). After digestion, an equal volume of ethyl acetate was added, the mixture was acidified to 0.5% TFA, vortexed, and phase-separated (15,800 g, 2 min). The aqueous phase was dried and reconstituted in 4% ACN, 0.1% TFA.

StageTips were prepared from CDS-SDB or AFS-SDB disks according to the published protocol^2)^. In this study, 200 µL pipette tips were used. SDB disks were punched with Hamilton 16-gauge (1.2 mm inner diameter). The following desalting steps were employed:

1. Washing: 80% ACN, 0.1% TFA, 2000 g × 3 min
2. Equilibration: 4% ACN, 0.1% TFA, 2000 g × 3 min
3. Sample loading: 4% ACN, 0.1% TFA, 1500 g × 3 min
4. Washing: 4% ACN, 0.1% TFA, 2000 g × 3 min
5. Elution: 80% ACN, 0.1% TFA, 1500 g × 3 min

Eluates were vacuum-dried and reconstituted in 4% ACN, 0.5% TFA prior to nanoLC/MS/MS. Sample loading per StageTip was 20 ng and 500 ng (each n = 3). Throughout this study, the sample loading amount into StageTips was referenced to the protein amount before the digestion process, as determined by BCA assay.

### 2.4. NanoLC/MS/MS analysis

Desalted peptides were analyzed on nanoLC/MS/MS with an Ultimate 3000 RSLCnano pump (Thermo Fisher Scientific, Waltham, MA, USA), an HTC-PAL autosampler (CTC Analytics, Zwingen, Switzerland) and a timsTOF HT mass spectrometer (Bruker, Bremen, Germany) operated in DDA-PASEF mode. Instrument settings were: TIMS ramp 100 ms; ion mobility 0.6–1.5 V s cm^−2^; one MS scan followed by 10 PASEF MS/MS; *m/z* 100–1700; a polygon filter to exclude singly charged ions; quadrupole isolation width *m/z* 2–3; and stepped collision energy along the mobility ramp (42/32/37/42/47/51 eV). For nanoLC, self-pulled needle columns (250 mm length, 100 μm ID, 6 μm needle opening) packed with Reprosil-Pur 120 C18-AQ 1.9 μm reversed-phase material (Dr. Maisch, Ammerbuch, Germany) were employed. The injection volume was 5 μL, and the flow rate was 500 nL min^−1^. Separation was achieved by applying a stepped linear gradient of 4–8% ACN (5 min), 8–32% ACN (60 min), 32–80% ACN (5 min), and 80% ACN (10 min) in 0.1% formic acid.

### 2.5. Data Analysis

Raw data were processed in FragPipe version 23.1. Peptides and proteins were identified through automated database searching using MSFragger version 4.3 and Philosopher version 5.1.2 against the human database (release 2023/05, 20,470 protein entries) from UniProtKB/Swiss-Prot. Mass tolerances for precursor and fragment ions were set to 20 ppm. Digestion mode was set Trypsin/P, allowing for up to two missed cleavages. Oxidation (M) and acetylation (protein N-term) were allowed as variable modifications. Carbamidomethylation (C) was set as a fixed modification. The false discovery rate (FDR) filter was set to 0.01 at both the peptide-spectrum match (PSM) and protein levels. For label-free quantification, IonQuant was used with the match-between-runs (MBR) option enabled (retention time tolerance 0.4 min, *m/z* tolerance 10 ppm, ion mobility (1/K0) tolerance 0.05), and the ion-level FDR threshold was set to 0.01.

### 2.6. Data availability statement

The MS raw data files and the analysis files containing the identification and quantification results for peptides and protein groups have been deposited at the ProteomeXchange Consortium (http://proteomecentral.proteomexchange.org) via the jPOST partner repository (https://jpostdb.org)^10)^ with the data set identifier JPST004241/PXD072126.

## 3. RESULTS AND DISCUSSION

Two commercially available SDB-embedded PTFE disks, CDS-SDB and AFS-SDB, were incorporated into StageTips following the standard fabrication protocol, using 1.2-mm punches to mount the disks inside 200-µL pipette tips (n = 3 each). Tryptic peptides derived from HeLa cell lysates (500 ng or 20 ng) were desalted using each StageTip, followed by nanoLC/MS/MS analysis and label-free quantification with match-between-runs. The results are summarized in Fig. 1.

**Figure 1.**
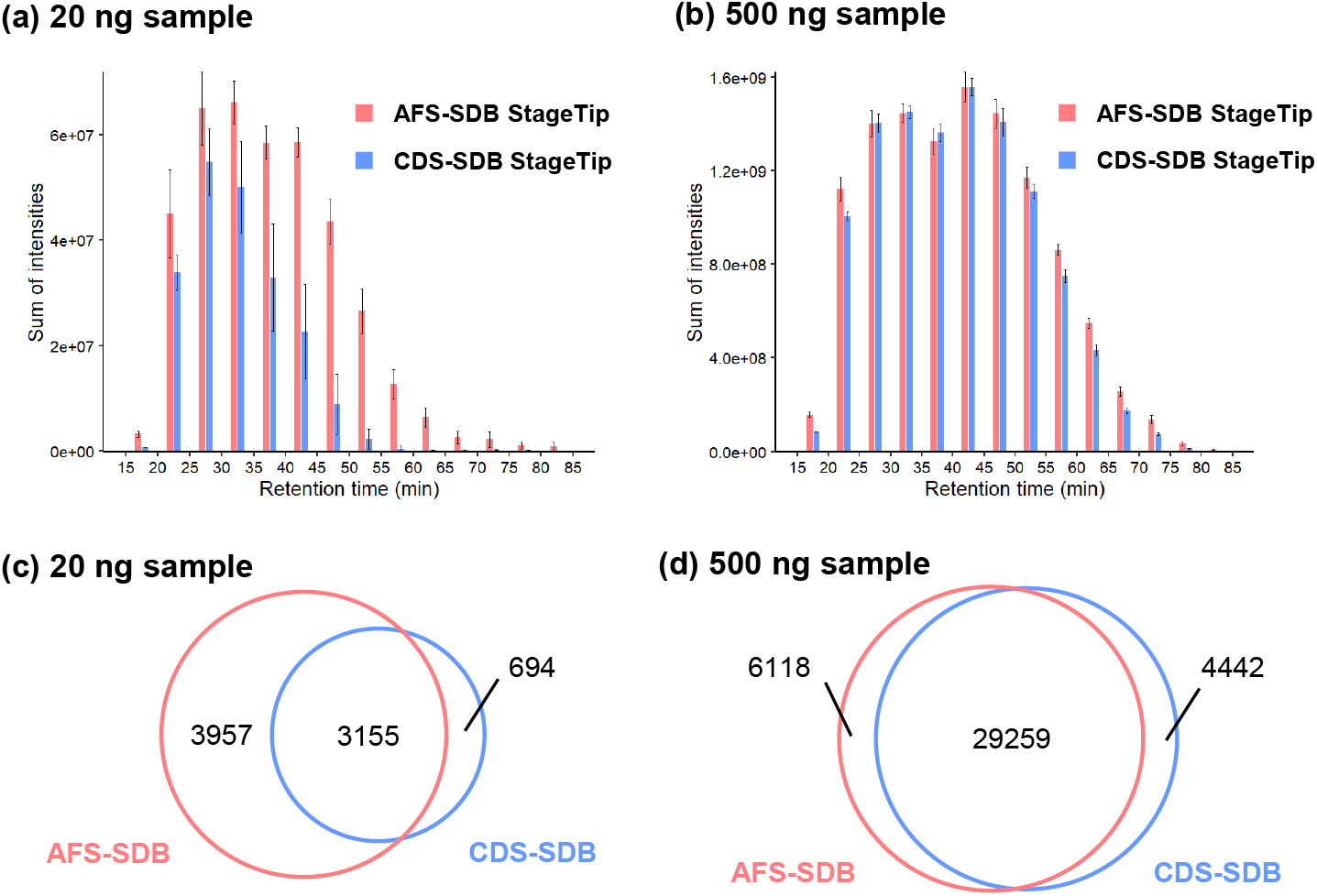
Comparison of AFS-SDB and CDS-SDB disks. (Upper panel) Comparison of the sum of identified peptide ion intensities binned by retention time between AFS-SDB and CDS-SDB StageTips. (a) 20 ng or (b) 500 ng of tryptic peptides from HeLa cell lysates were loaded onto StageTips. Fifty percent of the eluted peptides were injected into nanoLC/MS/MS. Error bars represent the standard deviation of triplicate analyses using three StageTips. (Lower panel) Overlap of identified peptides between AFS-SDB and CDS-SDB StageTips. (c) 20 ng or (d) 500 ng of tryptic peptides from HeLa cell lysates were loaded onto StageTips. Only peptides identified in all triplicate analysis were used for evaluation.

For the 500-ng samples, both disks yielded comparable peptide identifications. However, for the 20-ng samples, AFS-SDB identified 1.8-fold more peptides than CDS-SDB, with the largest differences observed among late-eluting, more hydrophobic peptides. Analysis of peptides uniquely identified by each disk revealed that AFS-SDB recovered significantly longer peptides (13.2 ± 4.4 amino acids) compared with CDS-SDB (9.6 ± 2.5 amino acids), suggesting reduced loss of long and hydrophobic peptides.

To investigate the basis of this difference, we compiled physical parameters from vendor documentation and measured disk dimensions and weight to estimate density (Table 1). AFS-SDB contains SDB particles reported to be 3.3 times larger, and the overall disk density was slightly lower (0.9 times). Considering that packing density does not depend on particle size, the AFS-SDB disk is thought to contain SDB particles packed somewhat more loosely. This is offset by the AFS-SDB disk being 40% thicker and containing 26% more stationary phase material than the CDS-SDB disk. In other words, these parameter comparisons alone cannot clearly explain the improved recovery of long peptides from low-input samples.

**Table 1.**
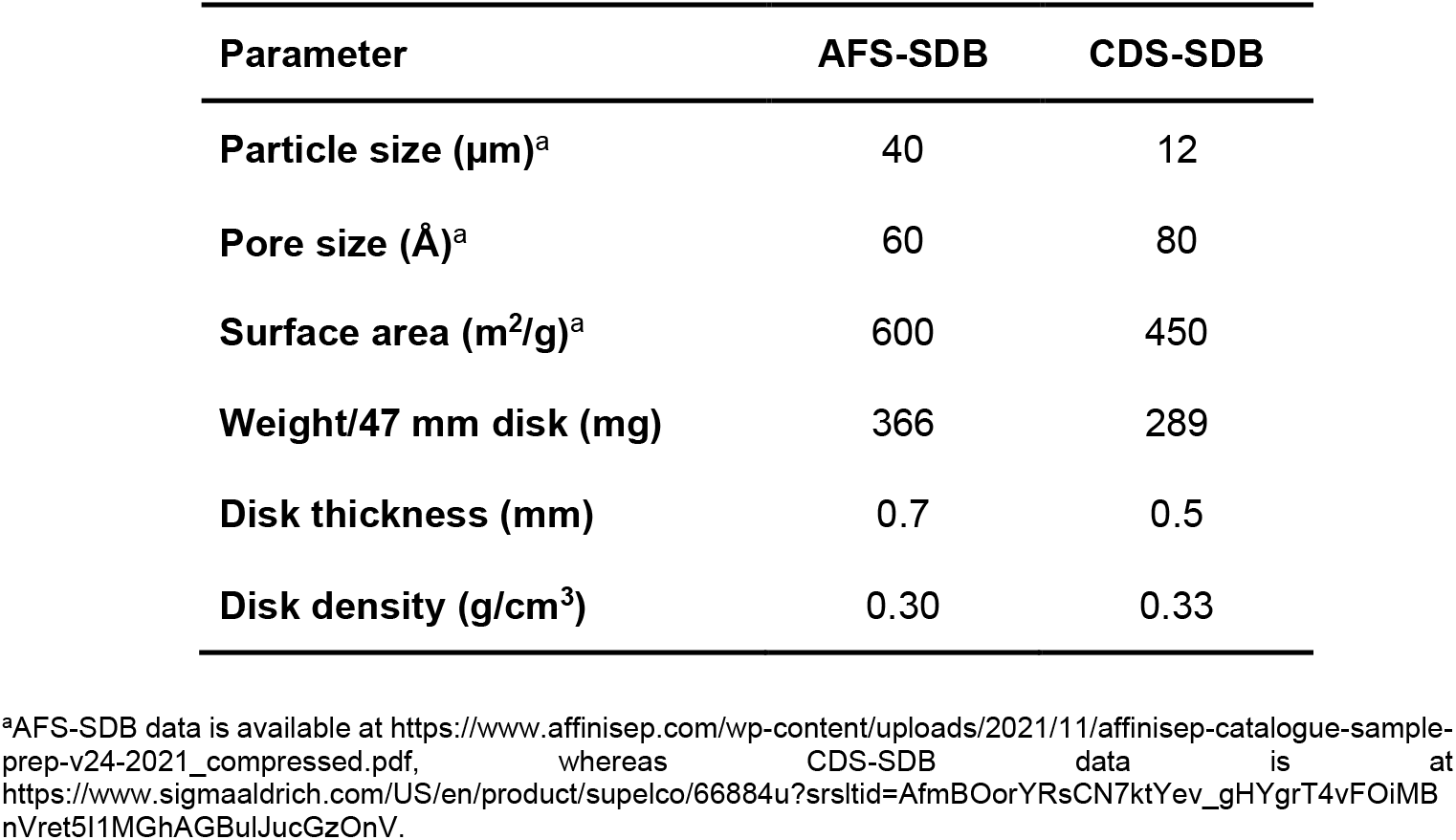
Physicochemical parameters of SDB disks.

The SEM imaging provided additional insight (Figure 2). While particle size of CDS-SDB is larger than reported (∼20 µm), the most striking distinction between two disks was the greater abundance of PTFE fibers in AFS-SDB. As shown in the enlarged image of Figure 2, these fibers adhere to the SDB particle surface and may partially cover the mesopores. A similar phenomenon was recently observed in our ChocoTip (CHrOmatographic particles COated by a Thermoplastic polymer Immobilized in Pipette Tip) disk, where interactions between the SDB particle surface and hydrophobic plastic polymers reduced the particle’s mesopores (10-100 nm), resulting in improved recovery of hydrophobic peptides, particularly at low sample amounts^11,12)^. In this case as well, the AFS-SDB disk containing more PTFE exhibited greater mesopore blockage. This likely inhibited the irreversible mesopore entry of long and hydrophobic peptides, thereby contributing to the improved recovery observed with AFS-SDB.

**Figure 2.**
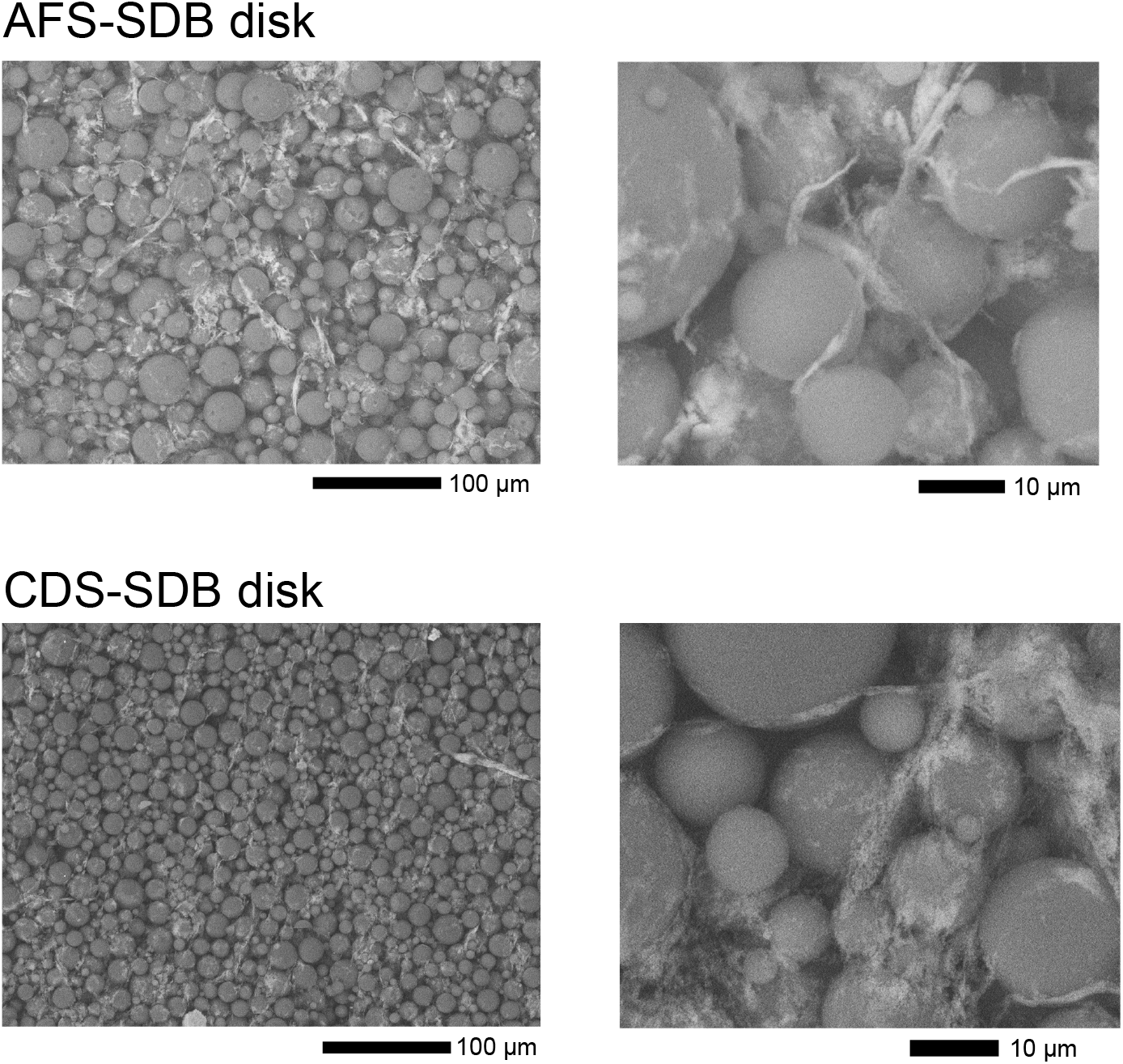
SEM imaging analysis of AFS-SDB and CDS-SDB disks. (Upper panel) AFS-SDB disk, (lower panel) CDS-SDB disk. Measurement parameters are described in the experimental section.

StageTips with commercially available disks employed in this study have inherent limitations in performance because they do not allow for customization of chromatography particle selection or SDB/PTFE ratios. In contrast, customizable StageTips like ChocoTip enable flexible control over the type and composition ratio of polymer fibers and chromatography particles^13)^, offering a promising pathway for developing StageTips optimized for trace amounts of samples including single-cell proteomics in future.

## ACKNOWLEDGMENTS

This work was funded by AMED-PRIME (24gm7010003h0001) and JST-ASTEP (JPMJTR25U9) to E. K., and JSPS Grants-in-Aid for Scientific Research 25K18607 to K. O. and 23H04924, 25K22526 to Y.I.

